## Supplementary information for "Haplotype Function Score improves biological interpretation and cross-ancestry polygenic prediction of human complex traits"

**Supplementary Notes**

**Simulation analysis**

We simulated trait levels using HFS data from chromosome 1 in a randomly selected 50,000 samples from UKB EUR training data. We randomly selected 1% (500) loci, assigned effect size from standard normal distribution, and calculated the aggregated genetic liability. We then simulated trait levels with h^2^=0.1. We applied HFS+SUSIE as well as REGENIE+PolyFun on simulated traits and calculated the Area Under Curve (AUC), false positive rate (FDR) at PIP>0.95 for HFS+SUSIE. We repeated this procedure for 30 times.

On average, HFS+SUSIE showed high accuracy in identifying causal loci (median AUC=0.92) and the FDR at PIP>0.95 is median 0.059. In line with real data analysis, the number of causal loci identified by HFS+SUSIE is 1.12-fold more than PolyFun on average. Furthermore, HFS+SUSIE showed good discrimination between causal and non-causal loci: the number of PIP>0.95 loci is larger than 0.5<PIP<0.95 loci.

**Alternative strategy on biological enrichment analysis**

Despite the standard linear regression as we applied in the main text, we also applied a linear mixed regression which took independent blocks as random effect. For each regression, we included one biological term plus all baseline annotations. The regression coefficient and p-value of each biological term were estimated by mgcv R package. After p-value correction, most of the significant terms were those recurrently appeared in more than half of the traits, which were considered artifacts of hidden covariates. When removing these recurrent terms, less than five significant terms remained for each trait. We concluded that linear mixed regression was less sensitive than standard linear regression for identifying trait-specific biological association.

We also tried another strategy for cell type-specific analysis. We first downloaded C8 category from MsigDB, which contained gene lists specifically expressed in about 800 cell types, derived from multiple single cell RNA sequencing studies. We then linked each locus to these gene lists by CS2G method, then applied linear regression, similar to pathway analysis. We found that most traits predominantly linked to nearly all cell types from a specific study, which showed study batch effect instead of biological functions. For example, smoking was associated with all neuron subtypes, pericytes and immune cells from one brain scRNA data, but did not showed association with immune cells and pericytes from other scRNA studies. We reasoned that the curated cell type-specific gene lists contained batch effects that were not yet corrected. Thus, in the main text, we reported association between PIP and single cell ATAC peak from one study, which reduced the batch effect.

**Hihglighted genes for complex traits**

For chronotype, we found one circadian gene CRY1 that were predicted to be target of locus chr12:1070930221107097118, which had PIP=0.56. This locus was active in cingulate gyrus, and belong to sequence class “enhancer-multi tissue”. CRY1 was known to participate in circadian pathway, and was not highlighted by previous GWAS. SNP-based fine-mapping also found no SNP with PIP>0.1 that was predicted to link to CRY1. We suggested that it was a novel promising target gene for understanding mechanisms of chronotype.

For systolic blood pressure, we found chr8:11726583-11730679 (PIP=0.999) that resided on gene GATA4. This locus was active in both adult heart ventricle and in fetal cardiomyocyte. GATA4 took part in physiological myocardial hypertrophy. SNP fine-mapping got PIP<0.34 for all SNPs linked to GATA4. Previous GWAS has found its homolog GATA2 as a key gene in blood pressure, and our new result supported GATA4 as another key genes.

For intelligence, we found chr10:45559452-45563548 that was active in caudate nucleus and was associated with intelligence at PIP>0.5. It was predicted to regulate ALOX5, a key enzyme in the Arachidonic acid metabolism. It is known that supplement of Arachidonic acid is beneficial for child intelligence development, and that arachidonic acid takes part in neurodevelopment. However, few genes related to arachidonic acid has been associated with intelligence.

**Figure S1 Distribution of number of haplotypes per locus**

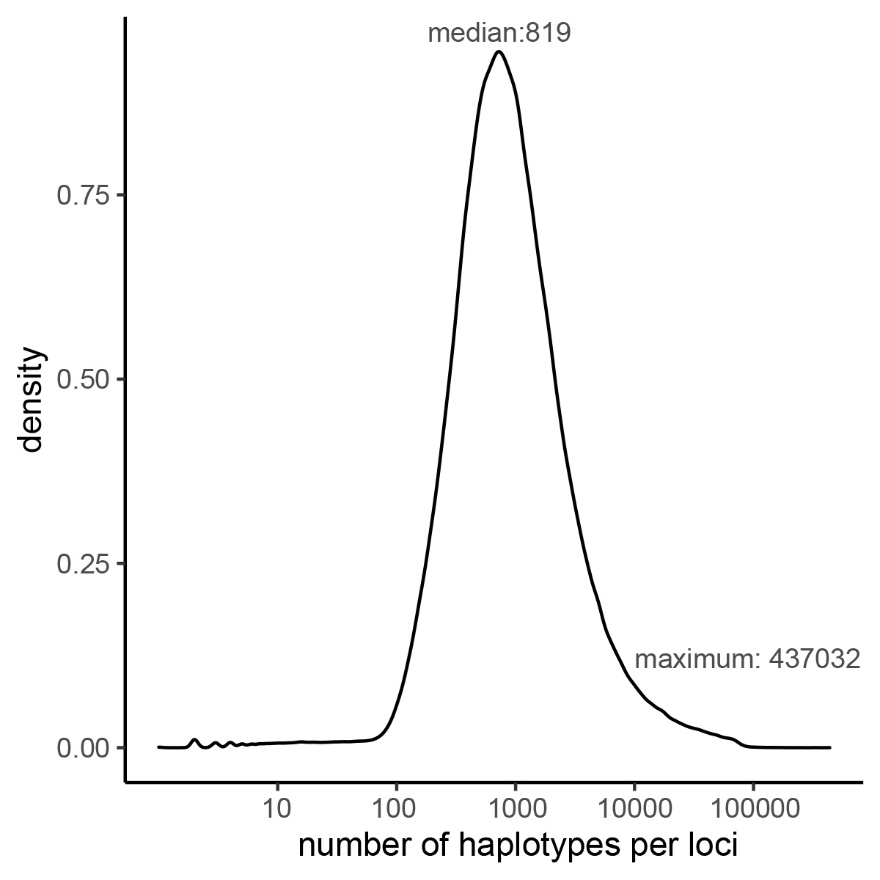

**Figure S2 Linkage disequilibrium among HFS.**

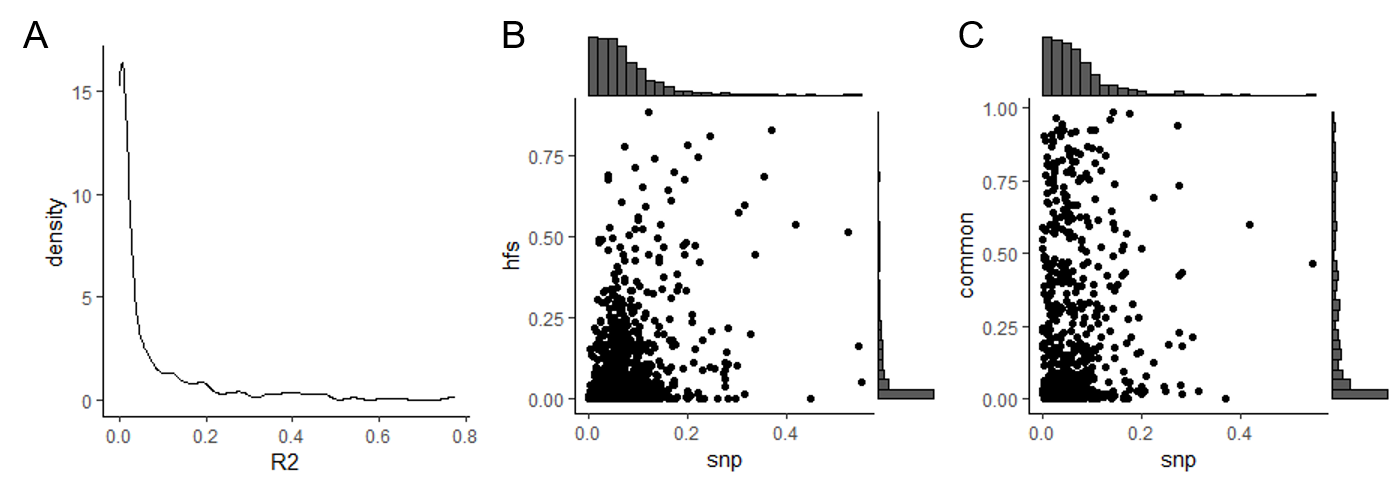

A: distribution of R^2^ of HFS from adjacent loci. B: comparison of R^2^ of adjacent loci HFS (y-axis) and median LD of SNP from the same adjacent loci (x-axis). C: same as B, but y-axis corresponded to HFS without rare variants.

**Figure S3 Comparison of inflation factor between HFS and SNP association tests**

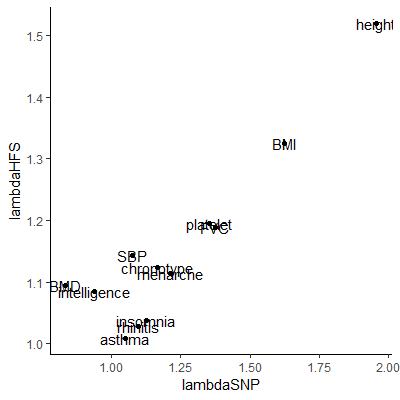

Lambda: genomic control inflation factors, defined as the median chi-squared statistics of all tested variables divided by 0.476.

**Figure S4 Heritability enrichment within causal loci estimated by LDSC**

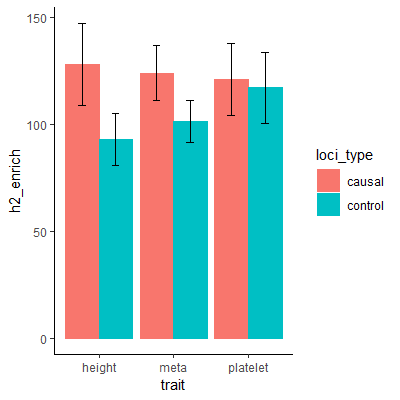

Causal loci: loci with PIP>0.95. Control loci: nearest locus of a causal locus that reached the same p-value level. R2_enrich: proportion of heritability divided by proportion of SNP. Meta: inverse variance-weighted heritability enrichment of test traits. Error bar indicated 95% confidence interval.

**Figure S5 P-value comparison within RBBP5 region between HFS and SNP association test**

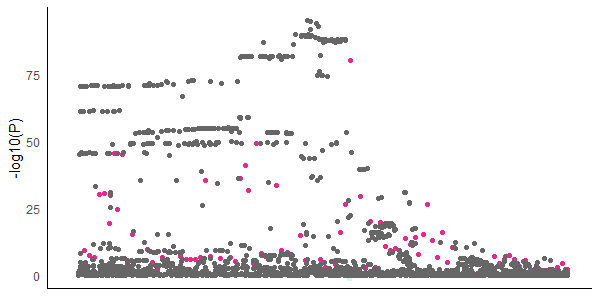

x-axis: Same chromosome region as Figure 4. Each black point represented a SNP, its y-axis represented REGENIE GWAS p-value. Each pink point represented a locus, its y-axis represented its p-value for HFS association test with platelet count.

**Figure S6 MHC locus analysis for allergic diseases.**

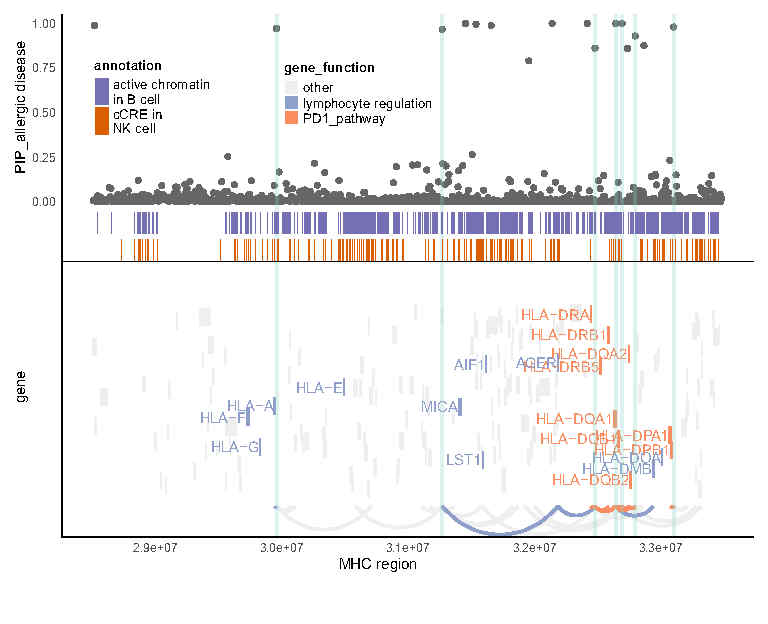

Similar to Figure 4C, but for other allergic diseases.

**Figure S7 Proportion of heritability captured by HFS PRS**

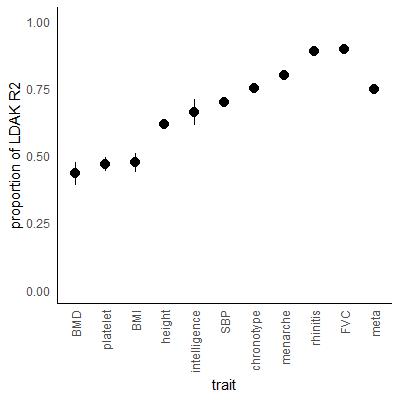

**Table S1 Basic information for genome-wide loci.**

Provided as separate .csv file.

**Table S2 Summary of HFS association tests for each trait.**

| trait | type | sample size | proportion of case | Nsig | Nind | lambdaSNP | lambdaHFS |
| --- | --- | --- | --- | --- | --- | --- | --- |
| height | continuous | 354072 | NA | 7573 | 1422 | 0.970582 | 0.753649 |
| BMI | continuous | 353708 | NA | 1482 | 387 | 0.804882 | 0.657047 |
| BMD | continuous | 205624 | NA | 1402 | 303 | 0.414095 | 0.543193 |
| platelet | continuous | 344284 | NA | 3102 | 619 | 0.67176 | 0.592315 |
| intelligence | continuous | 113441 | NA | 125 | 48 | 0.466088 | 0.538181 |
| SBP | continuous | 331497 | NA | 512 | 187 | 0.533585 | 0.566659 |
| chronotype | continuous | 316898 | NA | 240 | 76 | 0.578313 | 0.557171 |
| insomnia | continuous | 354592 | NA | 28 | 15 | 0.557961 | 0.514817 |
| menarche | continuous | 186261 | NA | 654 | 173 | 0.600759 | 0.552322 |
| FVC | continuous | 285641 | NA | 719 | 206 | 0.681845 | 0.589342 |
| asthma | binary | 350372 | 0.102976 | 355 | 80 | 0.52132 | 0.499852 |
| rhinitis | binary | 344258 | 0.204849 | 405 | 103 | 0.544623 | 0.5095 |

Nsig: number of loci that significantly associated with the trait. Nind: number of roughly independent loci associated with the trait. Lambda: genomic control inflation factor for HFS and SNP association tests.

**Table S3 Significant HFS-trait associations.**

Provided as separate .csv file.

**Table S4 Summary of HFS+SUSIE analysis with PIP>0.95.**

Provided as separate .csv file.

**Table S5 Summary of secondary analysis of HFS+SUSIE.**

Provided as separate .csv file.

**Table S6 Sequence class enrichment analysis for HFS+SUSIE result.**

Provided as separate .csv file. Source data of Figure 2.

**Table S7 Pathway enrichment result.**

Provided as separate .csv file.

**Table S8 Cell type and tissue enrichment result.**

Provided as separate .csv file.

**Table S9 Highlighted genes for platelet count.**

Provided as separate .csv file.

**Table S10 MHC region analysis result for asthma.**

Provided as separate .csv file.

**Table S11 MHC region analysis result for other allergic diseases.**

Provided as separate .csv file.

**Table S12 Highlighted genes for systolic blood pressure.**

Provided as separate .csv file.

**Table S13 Highlighted genes for chronotype.**

Provided as separate .csv file.

**Table S14 Highlighted genes for intelligence.**

Provided as separate .csv file.
